# A Novel Mistranslating tRNA Model in *Drosophila melanogaster* has Diverse, Sexually Dimorphic Effects

**DOI:** 10.1101/2021.09.17.460863

**Authors:** Joshua R. Isaacson, Matthew D. Berg, Jessica Jagiello, Judit Villén, Christopher J. Brandl, Amanda J. Moehring

## Abstract

Transfer RNAs (tRNAs) are the adaptor molecules required for reading of the genetic code and the accurate production of proteins. tRNA variants can lead to genome-wide mistranslation, the misincorporation of amino acids not specified by the standard genetic code into nascent proteins. While genome sequencing has identified putative mistranslating tRNA variants in human populations, little is known regarding how mistranslation affects multicellular organisms. Here, we create a *Drosophila melanogaster* model for mistranslation by integrating a serine tRNA variant that mistranslates serine for proline (tRNA^Ser^_UGG, G26A_) into the fly genome. Using mass spectrometry, we find that tRNA^Ser^_UGG, G26A_ misincorporates serine for proline at a frequency of ∼ 0.6% per codon. We find that mistranslation extends development time and decreases the number of flies that reach adulthood. Adult flies containing tRNA^Ser^_UGG, G26A_ present with more morphological deformities and worse climbing performance than flies expressing only wild type tRNA. Female flies with the serine tRNA variant have more deformities and experience a faster decline in climbing performance than males, suggesting sex-specific effects. This model will enable studies into the synergistic effects of mistranslating tRNA variants and disease-causing alleles.

## INTRODUCTION

Mistranslation occurs when an amino acid that differs from what is specified by the standard genetic code is incorporated into nascent proteins. Mistranslation disrupts proteostasis and impairs cell function and growth (Nangle *et al*. 2006; Paredes *et al*. 2012; Reverendo *et al*. 2014; Liu *et al*. 2014; Shcherbakov *et al*. 2019), yet naturally occurs in all cells at rates of 10^−2^ to 10^−5^ per codon, depending on the codon (reviewed in Joshi *et al*. 2019). Protein quality control pathways allow cells to tolerate mistranslation at frequencies approaching 10% in *Saccharomyces cerevisiae* and *Escherichia coli* (Ruan *et al*. 2008; Berg *et al*. 2019b). Mistranslating tRNA variants are also tolerated because of the buffering provided by multiple copies of the genes encoding tRNAs in most organisms (e.g. 610 total tRNA genes in humans and 295 total tRNA genes in *Drosophila melanogaster*, Chan and Lowe 2016). Mistranslation can be an adaptive response. For example, high levels of mistranslation increase survival of *E. coli* exposed to DNA damage (Samhita *et al*. 2020) and misincorporation of methionine protects mammalian, yeast, and bacterial cells against reactive oxygen species (Netzer *et al*. 2009; Wiltrout *et al*. 2012).

Mutations within tRNA encoding genes that alter specificity of aminoacylation or codon recognition increase mistranslation frequency. The fidelity of aminoacylation depends on aminoacyl-tRNA synthetases (aaRSs) correctly recognizing their cognate tRNAs through nucleotides, base pairs, and structural elements within the tRNA called identity elements. The anticodon is the main identity element for all tRNAs except tRNA^Ser^, tRNA^Ala^ and tRNA^Leu^ (Mcclain and Foss 1988; Hou and Schimmel 1988; Normanly *et al*. 1992; Asahara *et al*. 1993; Achsel and Gross 1993; Breitschopf *et al*. 1995; Himeno *et al*. 1997; Giegé *et al*. 1998). Changing the anticodon of tRNA^Ser^ does not affect aminoacylation but changes codon recognition (Garza *et al*. 1990; Geslain *et al*. 2010; Reverendo *et al*. 2014; Zimmerman *et al*. 2018; Berg *et al*. 2019b).

Mutations in tRNAs that cause mistranslation arise spontaneously and were identified initially in *E. coli* as suppressors of nonsense and missense mutations (see for examples; Stadler and Yanofsky 1959; Gorini and Beckwith 1966; Goodman *et al*. 1968). Subsequently, mistranslating tRNAs have been identified through their suppression of deleterious phenotypes in fungi, nematodes, plants, and mammalian cells (e.g Goodman *et al*. 1977; Wills *et al*. 1983; Chiu and Morris 1997; El Meziane *et al*. 1998; Murakami *et al*. 2005). While no spontaneous tRNA variants have been detected through suppression screens in *Drosophila*, Laski *et al*. (1989) and Garza *et al*. (1990) have engineered amber suppressing tRNA^Tyr^ and tRNA^Leu^ variants, respectively, that show a low level of amber stop codon suppression activity when integrated into the *Drosophila melanogaster* genome. In both cases sterility was noted.

In humans, mistranslation due to tRNA variants can cause disease (Goto *et al*. 1990; Shoffner *et al*. 1990; Zheng *et al*. 2012; Ishimura *et al*. 2014; Schoenmakers *et al*. 2016; reviewed in Kapur and Ackerman 2018 and Lant *et al*. 2019). Yet tRNA variants are relatively common in humans, with ∼66 tRNA variants per individual, some of which are predicted to decrease translational fidelity (Berg *et al*. 2019a). Despite the prevalence of potential mistranslating tRNAs and the potential links between mistranslation and disease, the impact of mistranslating cytoplasmic tRNAs in multicellular organisms is not well described. In this study, we develop a transgenic model of mistranslation in *D. melanogaster* by genomically integrating a serine tRNA variant that mistranslates serine at proline codons. We hypothesize that loss of proteostasis caused by the mistranslating tRNA will impact development, fitness, and behaviour. Serine for proline substitutions were detected by mass spectrometry in pupae expressing the mistranslating tRNA variant. Development time of flies containing the mistranslating tRNA was extended and fewer flies reached adulthood compared to wild type flies. The tRNA variant increased the prevalence of morphological deformities in adult flies, with females being more severely affected than males. Mistranslation also impaired climbing performance. These results demonstrate that *D. melanogaster* provides a model for the impact and genetic interactions of mistranslating tRNAs in a multicellular organism and their sexually dimorphic effects.

## METHODS

### Fly husbandry and stocks

All fly stocks were obtained from the Bloomington *Drosophila* Stock Centre and maintained on standard Bloomington recipe food medium (BDSC; Bloomington, Indiana) under a 14:10 light:dark cycle at 24°C and 70% relative humidity.

### Creating transgenic stocks

The gene encoding wild type tRNA^Ser^_UGA_ (FlyBase ID: FBgn0050201) was amplified from *D. melanogaster* genomic DNA using primers VK3400/VK3401 (primers are listed in Table S1) and cloned into pCDF4, which was a kind gift from Dr. Simon Bullock (Port *et al*. 2014) as a *Bgl*II/*Xba*I fragment to create pCB4222. The gene encoding a variant tRNA with a proline UGG anticodon and G26A secondary mutation (tRNA^Ser^_UGG, G26A_) were made by two step mutagenic PCR with primers VK3400/VK3889 and VK3401/VK3890 in the first round and pCB4222 as a template. Products from the first round were amplified with outside primers VK3400/VK3401 and cloned as a *Bgl*II/*Xba*I fragment into pCDF4 to give pCB4250. Full sequences of wild type tRNA^Ser^_UGA_ and tRNA^Ser^_UGG, G26A_ can be found in Figure S1.

To create flies containing mistranslating tRNAs, a stock expressing phiC31 (ΦC31) integrase in the germ line and containing an *attP* site in the left arm of the second chromosome was used (stock # 25709: *y*^*1*^ *v*^*1*^ *P{nos-phiC31\int*.*NLS}X; P{CaryP}attP40*). Plasmids were injected into *D. melanogaster* embryos using the protocol described in Isaacson (2018). Transgenic flies were identified by their wild type eye colour and balanced using stock # 3703 (*w*^*1118*^*/Dp(1;Y)y*^*+*^; *CyO/nub*^*1*^ *b*^*1*^ *sna Sco lt*^*1*^ *stw*^*3*^; *MKRS/TM6B, Tb*^*1*^) and #76359 (*w*^*1118*^; *wg*^*Sp-1*^*/CyO, P{w*^*+mC*^*=2xTb*^*1*^*-RFP}CyO; MKRS/TM6B, Tb*^*1*^) to create final stocks of the following genotype: *w*^*1118*^; *P{CaryP}attP40[v*^*+*^*=tRNA]*/*CyO, P{w*^*+*^*mC=2xTb*^*1*^*-RFP}CyO*; *MKRS*/*TM6B, Tb*^*1*^. After producing offspring, DNA was extracted from both parents of the final cross and PCR amplified using the primer set M13R and VK3400. PCR products were sequenced to confirm tRNA identity.

### Complementation in Saccharomyces cerevisiae

The *Bgl*II/*Xba*I fragment of pCB422 encoding *Drosophila* tRNA^Ser^_UGG, G26A_ was cloned into the *Bam*HI/*Xba*I sites of the yeast-*E. coli* shuttle plasmid YEPlac181 (Gietz and Sugino 1988; *LEU2*, 2 micron; CB4877). CB4877 and YEPlac181 were transformed into the yeast strain CY9013 (*MATα his3Δ1 leu2Δ0 lys2Δ0 met15Δ0 ura3Δ0 tti2Δ-met5Δ-mTn10luk* containing pRS313 (Sikorski and Hieter 1989) expressing *tti2-L187P* (Berg *et al*. 2017) selecting for growth on minimal plates lacking leucine and histidine. Transformants were streaked onto yeast-peptone (YP) plates containing 2% glucose and 5% ethanol and grown at 30°C for 4 days.

### Mass spectrometry

Six replicates of twenty pupae were collected from each genotype and lysed in 8 M urea, 50 mM Tris, 75 mM NaCl, pH 8.2 by grinding with a pestle and with glass beads at 4°C. Protein was reduced with 5 mM dithiothreitol for 30 minutes at 55°C and alkylated with 15 mM iodoacetamine for 30 minutes at room temperature. Robotic purification and digestion of proteins into peptides were performed on the KingFisher™ Flex using LysC and the R2-P1 method as described in Leutert *et al*. (2019). Peptides were separated by reverse-phase chromatography and online analyzed on a hybrid quadrupole orbitrap mass spectrometer (Orbitrap Exploris 480; Thermo Fisher Scientific) operated in data-dependent acquisition mode as described in Berg *et al*. (2021).

MS/MS spectra were searched against the *D. melanogaster* protein sequence database (downloaded from Uniprot in 2016) using Comet (release 2015.01; Eng *et al*. 2013). The precursor mass tolerance was set to 50 ppm. Constant modification of cysteine carbamidomethylation (57.0215 Da) and variable modification of methionine oxidation (15.9949 Da) and proline to serine (−10.0207 Da) were used for all searches. A maximum of two of each variable modification were allowed per peptide. Search results were filtered to a 1% false discovery rate at the peptide spectrum match level using Percolator (Käll *et al*. 2007). The mistranslation frequency was calculated using the unique mistranslated peptides for which the non-mistranslated sibling peptide was also observed. The frequency is defined as the counts of mistranslated peptides, where serine was inserted for proline, divided by the counts of all peptides containing proline, respectively, and expressed as a percentage.

### Scoring for deformities

Virgin, heterozygous flies were collected within ∼8 hours of eclosion and scored for deformities in adult legs (limbs gnarled or missing segments), wings (blistered, absent, fluid-filled, or abnormal size), or abdomen (fused or incomplete tergites). Flies collected before wing expansion were excluded. Sex and type of deformity was recorded. Flies that had multiple deformities had each recorded. 433 tRNA^Ser^_UGA_ flies (227 males and 216 females) and 656 tRNA^Ser^_UGG, G26A_ flies (345 male and 311 female) were scored. All deformities were photographed through the lens of a stereomicroscope using a Samsung Galaxy S8 camera.

### Developmental assays

Approximately 250 flies of each genotype were placed into fly cages and allowed to lay eggs for one hour. Three replicates of 30 eggs from each plate were collected and checked every 12 hours to record progress through each of the following developmental stages: egg hatching into larva, larva pupating into pupa, and adult eclosing from pupa. Sex, zygosity, and deformities (as described above) were recorded.

### Climbing assays

Virgin adult flies were collected, sorted by sex, and scored for deformities. Deformed flies or flies homozygous for the transgenic tRNA were discarded. Equal numbers of flies were collected from each genotype during each collection period. Sixty flies in 11 vials from each genotype were collected and transferred to new food the day before testing. The number of flies that climbed to a 5cm line in 10 seconds was recorded, and flies were retested every three days until the flies were 51 days old. Each vial was tested three times.

### Statistical analyses

All statistical analyses were performed using R Studio 1.2.5001. Frequency of proline-to-serine misincorporation between tRNA^Ser^_UGA_ and tRNA^Ser^_UGG, G26A_ was compared using a *t*-test. Developmental time data were compared using Wilcoxon rank-sum tests. Fisher’s exact tests were used to compare survival between developmental stages and proportion of deformities. Chi-square tests were used to compare prevalence of each type of deformity using a post hoc analysis outlined in Shan and Gerstenberger (2017). A generalized linear model was constructed from the climbing assay data and performance was compared using F-tests.

## RESULTS

### *A tRNA*^*Ser*^ *variant induces mistranslation in* Drosophila melanogaster

To characterize mistranslation in a multicellular organism, we integrated genes encoding wild type tRNA^Ser^_UGA_ as a control and a tRNA^Ser^ variant that mistranslates serine for proline (Figure 1A) into the left arm of the second chromosome of the *D. melanogaster* genome. The tRNA^Ser^ variant has a proline UGG anticodon and G26A secondary mutation (tRNA^Ser^_UGG, G26A_). The alleles were balanced over a homolog that has serial inversions, preventing recombinant offspring and transgene loss. tRNA insertions were validated with PCR using primers specific to the inserted plasmid and confirmed by sequencing. The secondary G26A mutation was included in the mistranslating tRNA to dampen tRNA function as we have previously found a tRNA^Ser^ variant with a proline anticodon causes lethal levels of mistranslation when expressed in yeast (Berg *et al*. 2017).

**Figure 1.**
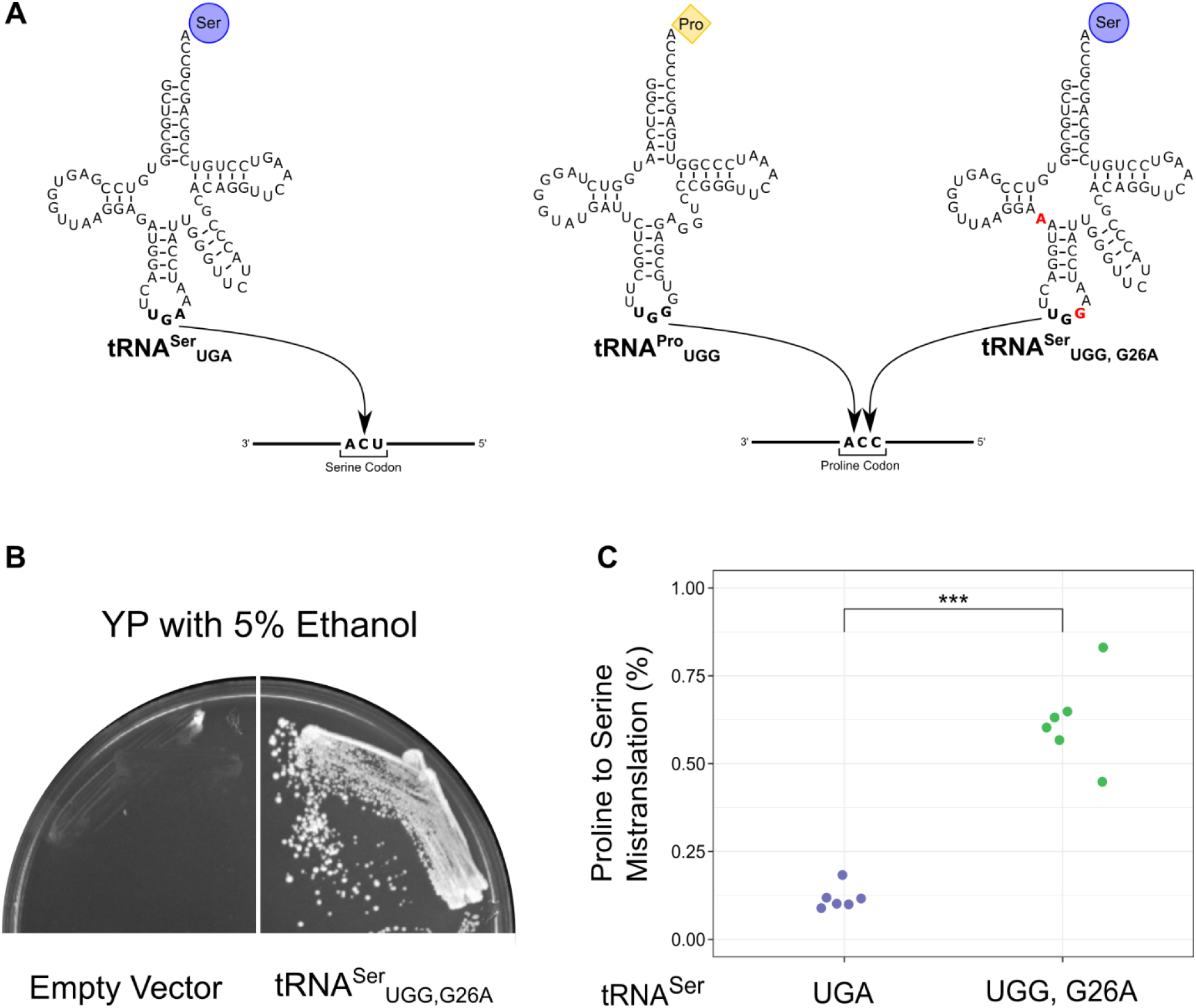
tRNA^Ser^_UGG, G26A_ induces mistranslation in D. melanogaster. **A)** Wild type serine tRNAs base pair with serine codons during translation and incorporate serine into the growing polypeptide. Serine tRNA variants with anticodon mutations can recognize non-serine codons and misincorporate serine during translation. tRNA^Ser^_UGG, G26A_ competes with tRNA^Pro^_UGG_ for CCA codons and inserts serine at proline codons. Red bases indicate mutation compared to the wild type tRNA^Ser^_UGA_. **B)** *D. melanogaster* tRNA^Ser^_UGG, G26A_ suppresses the ethanol sensitivity caused by *tti2-L187P* in *S. cerevisiae*. Plasmid encoding the vector alone (left) or the gene expressing tRNA^Ser^_UGG, G26A_ (right) were transformed into CY9013 (*tti2-L187P*), streaked onto YP medium containing 5% ethanol and grown at 30° for 4 days. **C)** Frequency of proline-to-serine mistranslation in tRNA^Ser^_UGA_ and tRNA^Ser^_UGG, G26A_ pupae (n = 6 replicates of 20 pupae each). Genotypes were compared using a *t*-test. “***” p < 0.001.

Adults homozygous for tRNA^Ser^_UGA_ or tRNA^Ser^_UGG, G26A_ can be produced. However, we were unable to propagate the strain homozygous for tRNA^Ser^_UGG, G26A_ because crosses between male and female tRNA^Ser^_UGG, G26A_ homozygotes produce no viable offspring. As such, we used heterozygous flies for our experiments with adults. Studying heterozygous flies may be more biologically relevant as mistranslating tRNAs present in populations are likely to arise as single alleles. We determined zygosity by balancing the tRNAs over a *CyO* homolog containing Tubby-linked RFP and *miniwhite* (Pina and Pignoni 2012). Heterozygous larvae and pupae are identified by the presence of RFP and heterozygous adults by their curly wings and non-white eyes.

As an initial test of mistranslation by *Drosophila* tRNA^Ser^_UGG, G26A_, we determined if it would rescue the growth of a *S. cerevisiae* strain containing *tti2-L187* (CY4013). The *tti2-L187* allele contains a missense mutation converting a CUA codon for leucine to CCA for proline and results in the slow growth of yeast in stress conditions including in medium containing 5% ethanol (Hoffman *et al*. 2017). Mistranslation of proline to serine rescues the growth of yeast cells in ethanol medium (Berg *et al*. 2017). The gene encoding *Drosophila* tRNA^Ser^_UGG,G26A_ was transformed into yeast strain CY4013 that contains *tti2-L187* as the sole copy of *TTI2*. Cells were transformed with plasmid expressing *Drosophila* tRNA^Ser^_UGG,G26A_ or vector alone. As shown in Figure 1B, *Drosophila* tRNA^Ser^_UGG,G26A_ enabled growth of CY4013 on medium containing 5% ethanol indicative of mistranslation by *Drosophila* tRNA^Ser^_UGG, G26A_.

We then analyzed the proteome of *D. melanogaster* pupae by mass spectrometry to determine the mistranslation frequency (Figure 1C; Supplemental File S2). Pupae were used because of the extensive cellular remodelling and corresponding rapid changes in protein synthesis that occur during this stage (Mitchell *et al*. 1977; Mitchell and Petersen 1981), and the potential of mistranslation during this stage to influence adult traits such as anatomy or neuronal function. The frequency of proline to serine mistranslation calculated as the ratio of peptides containing the mistranslated serine residue to peptides containing the cognate proline residue was ∼0.6% in flies expressing tRNA^Ser^_UGG, G26A_. In the control strain, the frequency of proline to serine substitutions was 0.1%.

### Mistranslation adversely affects D. melanogaster development

To determine if tRNA^Ser^_UGG, G26A_ affects fly development, we collected 90 one-hour old embryos and monitored how many individuals survived through each developmental stage and time between each stage: egg laying to embryos hatching into larvae, hatching to pupation, and pupation to eclosion of adults. Since the RFP marker used to determine tRNA zygosity is not expressed at early embryonic stages, both homozygotes and heterozygotes were pooled in this assay. Flies were checked every twelve hours and sex, zygosity, and presence of deformities in adults were recorded (Supplemental file S2). No obvious patterns were noted regarding these traits and because too few adults eclosed in this experiment, statistical comparisons were not possible.

Of the 90 embryos collected, significantly more flies with wild type tRNA^Ser^_UGA_ reached larval (41 vs. 27, Fisher’s exact test, p = 0.045), pupal (24 vs. 10, p = 0.013), and adult stages (22 vs. 9, p = 0.017, Figure 2A) than the mistranslating tRNA^Ser^_UGG, G26A_. Absolute numbers are influenced by die-off in the preceding life-cycle stage, so proportions were also calculated to assess pupal and adult survival. When comparing the proportion of larvae that pupated, 58% of tRNA^Ser^_UGA_ and 37% of tRNA^Ser^_UGG, G26A_ larvae reached pupation, although this difference was not statistically significant (p = 0.13). The proportion of adults that eclosed from pupae was virtually identical, 91% for tRNA^Ser^_UGA_ and 90% for tRNA^Ser^_UGG, G26A_ (Supp. File S2). These data indicate that flies are particularly susceptible to lethal effects of mistranslation during embryogenesis but show increased tolerance once they reach pupation. Although there was no significant difference between proportion of tRNA^Ser^_UGA_ and tRNA^Ser^_UGG, G26A_ larvae that pupated, there was a trend of lower survival in tRNA^Ser^_UGG, G26A_ such that larvae seem to display an intermediate sensitivity phenotype between embryos and pupae.

**Figure 2.**
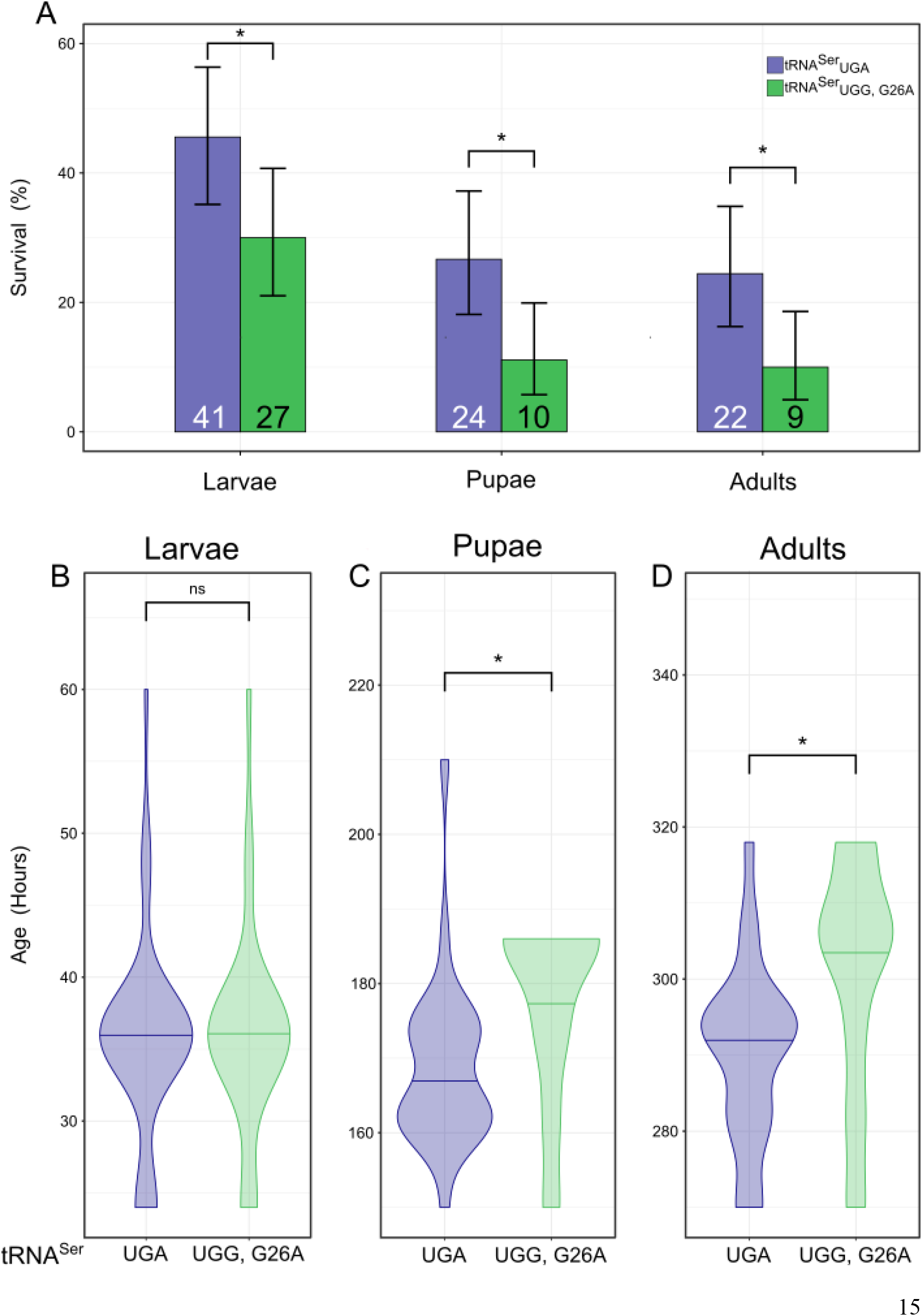
Mistranslation from a tRNA variant impacts development of D. melanogaster. **A)** Percentage of the 90 tRNA^Ser^_UGA_ and tRNA^Ser^_UGG, G26A_ containing individuals that made it to larval, pupal, and adult stages. Survival was compared using Fisher’s exact test. Error bars represent the 95% confidence interval of the proportion. Values within bars represent the number of flies that reached that developmental stage. **B)** Violin plot depicting the distribution of times until for tRNA^Ser^_UGA_ and tRNA^Ser^_UGG, G26A_ embryos to hatch into larva. The horizontal line within the plot represents the median of the distribution. Genotypes were compared using Wilcoxon rank-sum tests. **C)** Total time until pupation. **D)** Total time until eclosion. “ns” p ≥ 0.05, “*” p < 0.05, “**” p < 0.01, “***” p < 0.001.

Eggs expressing tRNA^Ser^_UGG, G26A_ had similar hatching times as eggs expressing wild type tRNA^Ser^ (p = 0.78, Wilcoxon rank-sum test, Figure 2B). However, larvae expressing tRNA^Ser^_UGG, G26A_ pupated significantly slower than the wild type (p = 0.023, Figure 2C). This trend continued into adulthood, as the control adult tRNA^Ser^_UGA_ flies eclosed significantly sooner than tRNA^Ser^_UGG, G26A_ flies (p = 0.047, Figure 2D). Only 20% of the mistranslating flies pupated by the median time for flies with the wild-type tRNA, and only 33% eclosed by the wild-type tRNA median

Mutations in genes vital to proteostasis or translation fidelity cause morphological defects (Rutherford and Lindquist 1998; Cui and DiMario 2007; Reverendo *et al*. 2014). We observed that flies containing one copy of the exogenous tRNA^Ser^_UGG, G26A_ had deformities including gnarled or blistered legs, notched wings, and misfused tergites (Figure 3A-D). Other abnormalities (e.g. haltere aberrations or rough eyes) were rarely observed, so only the more common leg, wing, and tergite deformities were scored. To determine if the frequency of deformities was greater than the control, we calculated the proportion of flies that eclosed with at least one deformity. These flies were collected separately from the development assay described above. From a total of 433 tRNA^Ser^_UGA_ flies (227 males and 216 females) and 656 tRNA^Ser^_UGG, G26A_ flies (345 male and 311 female) we identified significantly more deformities in flies containing tRNA^Ser^_UGG, G26A_ than tRNA^Ser^_UGA_ (Fisher’s exact test corrected using Holm-Bonferroni’s method, p < 0.001, Figure 3E). In addition, female flies containing tRNA^Ser^_UGG, G26A_ had more deformities than males (p < 0.001, Figure 3F). Interestingly, flies containing tRNA^Ser^_UGG, G26A_ presented with disproportionately more tergite deformities than flies with the wild type tRNA^Ser^_UGA_ (Chi-square test, corrected p = 0.03, Supplemental File S3), indicating that this mistranslating tRNA^Ser^ variant is particularly deleterious to fly abdominal development. These results suggest that mistranslation disrupts fly development and female flies are more sensitive to the presence of mistranslating tRNA variants.

**Figure 3.**
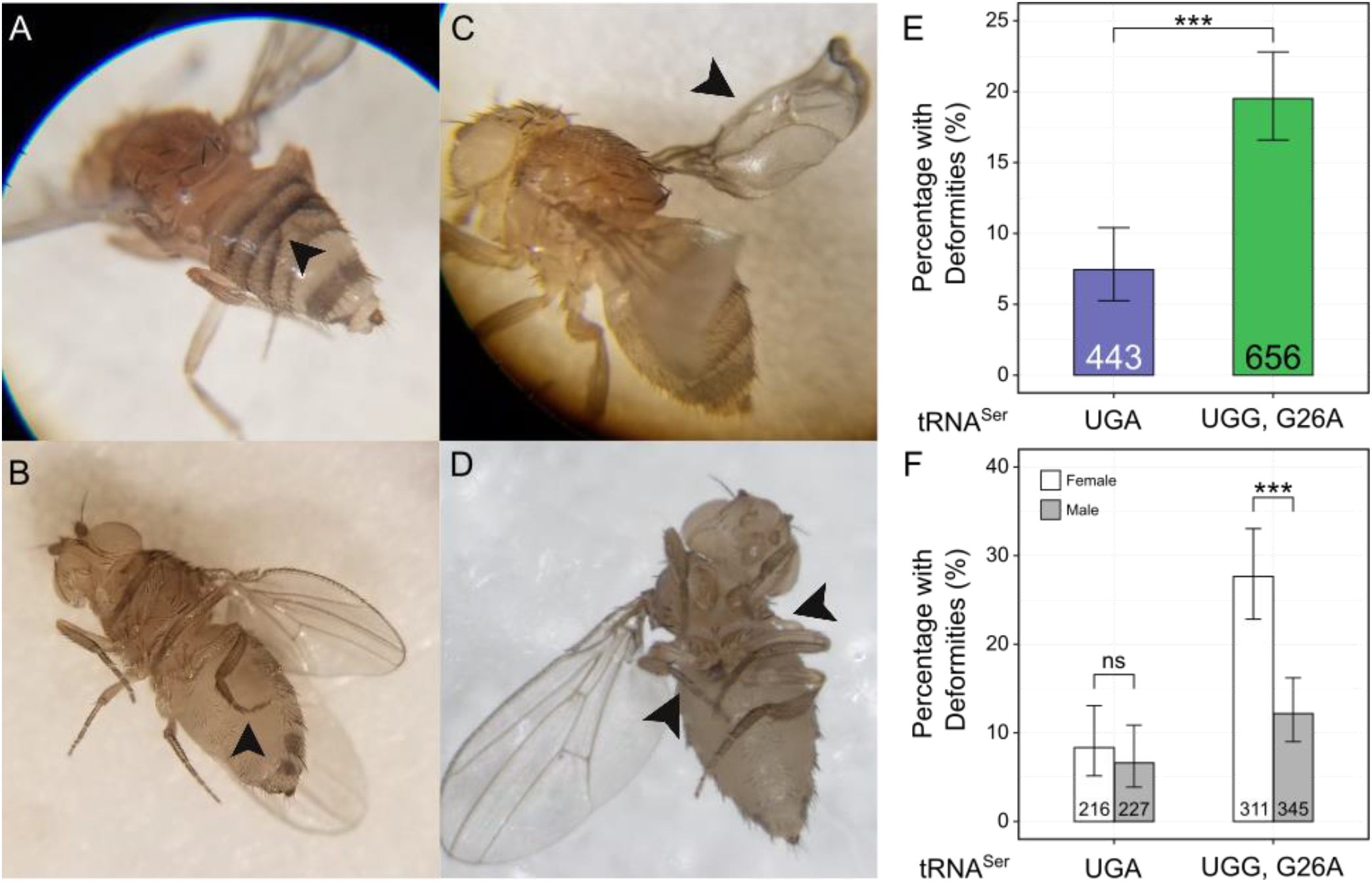
The tRNA^Ser^_UGG, G26A_ variant causes morphological deformities in adults in a sex-specific manner. **A)** Examples of flies with misfused tergites, **B)** gnarled hindlegs, **C)** wing blisters, and **D)** missing wings/legs, as indicated by arrowheads. **E)** Percentage of tRNA^Ser^_UGA_ or tRNA^Ser^_UGG, G26A_ flies that eclosed with at least one deformity. Groups were compared using Fisher’s exact test and corrected using the Holm-Bonferroni method. Bar height represents the percentage of flies of a genotype that had at least one deformity. Error bars represent the 95% confidence interval. Values within bars describe the number of flies examined for deformities. **F)** Same data as **E** but separated by sex. “ns” p ≥ 0.05, “*” p < 0.05, “**” p < 0.01, “***” p < 0.001.

### tRNA^Ser^_UGG, G26A_ impacts fly motility

Negative geotaxis assays are often used to study neurodegenerative diseases (e.g Feany and Bender 2000; Song *et al*. 2017; Aggarwal *et al*. 2019), so as an initial examination of neurodegeneration, we determined if tRNA^Ser^_UGG, G26A_ impaired climbing performance. Sixty virgin, heterozygous flies of the four genotypes (tRNA^Ser^_UGA_ males and females, and tRNA^Ser^_UGG, G26A_ males and females) were collected and tested using a climbing assay every three days; flies with deformities were not used in this experiment.

As expected, climbing performance of all genotypes decreased with age (F-tests performed on generalized linear models corrected using Bonferroni’s method, Supplementary File S2). For both males and females, climbing performance of tRNA^Ser^_UGG, G26A_ flies was significantly worse than wild type tRNA^Ser^_UGA_ flies (male: p = 0.001, female: p < 0.001, Figure 4A, B). Climbing performance was not significantly different when comparing males to females in either the control tRNA^Ser^_UGA_ (p = 0.08) or mistranslating tRNA^Ser^_UGG, G26A_ flies (corrected p → 1, Figure 4C, D). The climbing ability of male and female flies containing the wild type tRNA^Ser^_UGA_ declined at similar rates, as evidenced by the parallel performance curves (p = 0.97, Figure 4C). However, the climbing performance curve of female flies containing tRNA^Ser^_UGG, G26A_ intersected the male curve, indicating that female climbing performance declined faster than in males (p = 0.038, Figure 4D, Supp. File S3). These data suggest that the mistranslating tRNA^Ser^ variant negatively affects locomotion and has an accelerated impact on female ability to climb as they age.

**Figure 4.**
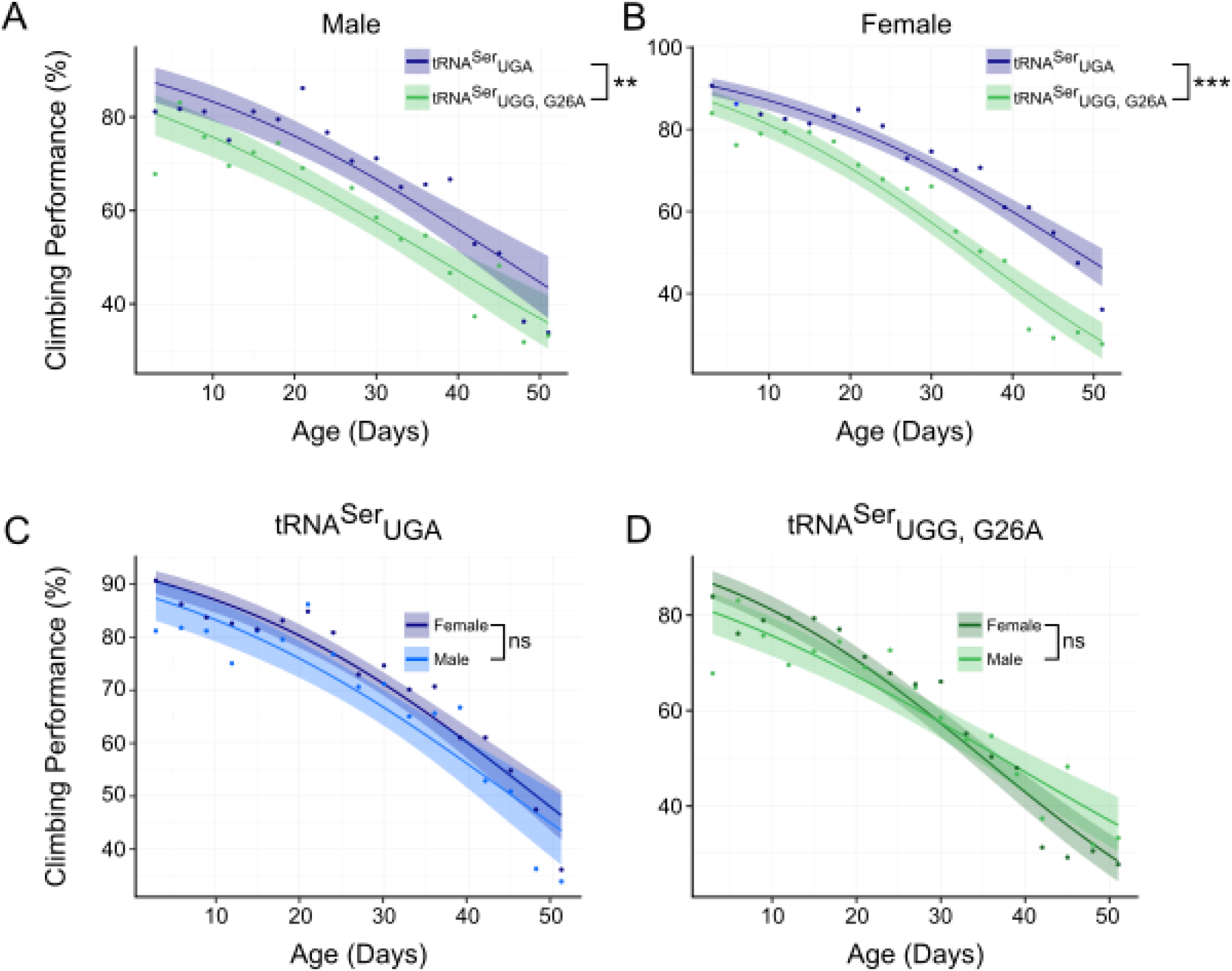
Fly locomotion is impacted by a mistranslating tRNA^Ser^ variant. Each point represents the percentage of flies (out of 60 from 11 vials) that managed to climb 5cm in ten seconds averaged over three trials. Generalized linear models were constructed from the performance data and F-tests were performed on the models. P-values were corrected using Bonferroni’s method. Shaded region represents the 95% confidence intervals for the fitted performance curves. **A)** Climbing performance of male flies containing tRNA^Ser^_UGA_ or tRNA^Ser^_UGG, G26A_. **B)** Climbing performance of female flies containing tRNA^Ser^_UGA_ or tRNA^Ser^_UGG, G26A_. **C)** Climbing performance of male and female flies containing tRNA^Ser^_UGA_ or **D)** tRNA^Ser^_UGG,G26A_. “ns” p ≥ 0.05, “*” p < 0.05, “**” p < 0.01, “***” p < 0.001.

## DISCUSSION

### Creating a fly model of mistranslation

We have created a *Drosophila melanogaster* model containing a genomically-integrated cytosolic tRNA that mistranslates serine for proline. The mistranslating fly model expands the possibilities offered by yeast or cell lines by allowing for studies into sex-specific or tissue-specific effects of mistranslating tRNA variants and the effect of tRNA variants on development and disease.

Our method of transgene integration controlled for positional effects by inserting either wild type or mistranslating tRNA^Ser^_UGG, G26A_ into the same locus on chromosome 2L. The fly lines containing tRNA^Ser^_UGG, G26A_ have not lost the transgene for over two years, indicating that mistranslating tRNA variants can be stably maintained in the genome. We observed a proline-to-serine misincorporation rate of ∼ 0.6% in the pupae for the genomically integrated tRNA^Ser^_UGG, G26A_ gene. This level of mistranslation was sufficient to cause deleterious phenotypes affecting diverse aspects of fly physiology.

Mistranslation frequency is affected by multiple factors, such as codon usage, expression of the tRNA, stability and turnover of the tRNA variant, and number of competing tRNA genes. The distribution of tRNAs that compete with tRNA^Ser^_UGG, G26A_ in flies as well as the relevant codon usage is shown in Supplemental File 1. Flies contain two proline tRNA species that compete with tRNA^Ser^_UGG, G26A_ for decoding CCA codons (tRNA^Pro^_UGG_ and tRNA^Pro^_AGG_ through modification of A34 to inosine; Crick 1966; Boccaletto *et al*. 2018); likewise mistranslating tRNA^Ser^_UGG, G26A_ competes with two proline tRNA species (tRNA^Pro^_CGG_ and tRNA^Pro^_UGG_ through wobble pairing) for decoding CCG codons. The maximum frequency of mistranslation with a heterozygous tRNA^Ser^_UGG, G26A_ encoding gene in flies is 2.4% (see Table S2 for calculations). The less than maximal frequency of 0.6% observed for tRNA^Ser^_UGG, G26A_ is expected because the G26A mutation prevents dimethylation at position 26, which increases turnover by the rapid tRNA decay pathway (Dewe *et al*. 2012).

### A mistranslating tRNA^Ser^ variant has diverse and sex-specific effects on flies

The mistranslating tRNA^Ser^_UGG, G26A_ affected fly physiology consistent with organism-wide loss of proteostasis. Our findings resemble other studies of proteostasis loss in flies. Impaired heat shock response exacerbates neurodegeneration and increases development time (Warrick *et al*. 1999; Gong and Golic 2006), and many of the wing, leg and tergite deformities observed for heterozygous *Heat shock protein 83* (*Hsp83*) mutants look similar to those observed in this study (Rutherford and Lindquist 1998). Developmental and neurodegenerative phenotypes including locomotive defects as measured in a climbing assay were likewise observed in flies containing a misacylation-prone PheRS (Lu *et al*. 2014). It is interesting to note that reduced levels of translation lead to similar deformities as found in mistranslating flies. RNAi knockdown of *Nopp140*, a gene involved in ribosome assembly, causes flies to present with gnarled legs, missing wings, and misfused tergites (Cui and DiMario 2007). *Minute* genes describe a collection of >50 genes required for protein synthesis. Their mutation results in shorter, thinner bristles, delayed development, smaller body size, and anatomical deformities when mutated (Schultz 1929; Marygold *et al*. 2007), again similar to the developmental and anatomical aberrations seen in flies containing the mistranslating tRNA^Ser^ variant. Though reduced translation and mistranslation are different processes, the similar phenotypes produced demonstrate that development is highly dependent on accurate and efficient translation.

The increased impact of the mistranslating tRNA on female flies was striking. *D. melanogaster* males and females have highly different physiology and experience different developmental challenges. Adult females are larger than males, develop faster, invest more resources into reproduction, and tend to live longer than males (Bonnier 1926; Bakker 1959; Sørensen *et al*. 2007; Ziehm *et al*. 2013). Males and females also display dimorphic responses to proteotoxic stress. Fredriksson *et al*. (2012) examined protein carbonylation in female somatic and germ line cells at different ages to determine how aging affects protein quality control of somatic and reproductive tissues. They found that as females age, there are fewer carbonylated proteins and reduced protein aggregation (both indicators of proteostasis loss) in eggs compared to the soma. Their work shows that females prioritize proteostasis of their eggs over their somatic cells, even while unmated. This trade-off could exacerbate the stress of mistranslating tRNAs in females, particularly as they experience aging-induced loss of proteostasis, and could contribute to the faster decline of climbing performance observed in female tRNA^Ser^_UGG, G26A_ flies compared to males. Many stress-response pathways affect males and females differently. For example, induction of the heat shock response increases male lifespan whereas female lifespan is unaffected (Sørensen *et al*. 2007; reviewed in Tower 2011). Dietary restriction shows the opposite trend, as it increases female lifespan more than male (Nakagawa *et al*. 2012; Regan *et al*. 2016; reviewed in Garratt 2020). Experiments testing the effects of mistranslating tRNAs on male and female fly longevity are ongoing. It is also possible that expression of the mistranslating tRNA differs between males and females or that the mistranslating tRNA has alternative functions (e.g. tRNA-derived fragments) that differ between males and females.

### Implications for human disease

Our work suggests that mistranslating tRNA variants have the potential to influence multiple aspects of human physiology. From a development perspective, the alteration in progression through life stages and increased number of deformities suggest that the proteotoxic stress resulting from mistranslating tRNA variants may contribute to congenital or developmental anomalies. Flies expressing tRNA^Ser^_UGG, G26A_ have a pattern of locomotion defects similar to those seen for flies expressing alleles associated with neurodegeneration (Feany and Bender 2000; Song *et al*. 2017; Aggarwal *et al*. 2019). Interestingly, the mistranslating fly model further resembles human neuropathies in that climbing performance declined faster in female compared to male flies, just as some neurodegenerative disorders such as Alzheimer’s and Huntington’s Disease, are more common or severe in women compared to men (Viña and Lloret 2010; Zielonka *et al*. 2013).

Given the prevalence of putative mistranslating tRNAs in the human population (Berg *et al*. 2019a) and the potential for mistranslation to disrupt proteostasis, we hypothesize that mistranslating tRNAs can exacerbate diseases characterized by a loss of proteostasis (see also Reverendo *et al*. 2014), and our results here indicate that these effects may differ in magnitude between sexes. Our previous studies in yeast have shown negative genetic interactions between mistranslation and mutations in genes involved in protein quality control and other pathways that could contribute to disease (Hoffman *et al*. 2017; Berg *et al*. 2020, 2021). Our *D. melanogaster* model of mistranslation allows for the expansion of these studies into the investigation of mutant tRNA contribution to disease and development.

### Data availability

Fly lines and plasmids are available upon request. The authors affirm that all data necessary for confirming the conclusions of the article are present within the article, figures, and supplemental material. Supplemental files are available at FigShare. Supplemental File S1 contains all supplemental figures and tables. Supplemental File S2 contains all raw data. Supplemental File S3 contains R code used to analyze mass spectrometry, developmental, deformity, and climbing assay data. Supplemental File S4 contains all images of deformed flies. The mass spectrometry proteomics data have been deposited to the ProteomeXchange Consortium via the PRIDE partner repository (Perez-Riverol *et al*. 2019) with the dataset identifier PXD028498.

## Acknowledgements

We would like to thank the Biotron Integrated Microscopy facility for their aid with this research. In particular, JRI would like to thank Karen Nygard, Reza Khazaee, and Marc Courchesne for their instruction and help. We would also like to thank Dr. Yolanda Morbey for guidance on analyzing the climbing assay data, Ricard Rodriguez-Mias for assisting with the mass spectrometry and maintaining the instruments, and Julie Genereaux for technical assistance.

## Funding

This work was supported by NSERC grants to CJB [RGPIN-2015-04394] and AJM [RGPIN-2020-06464], as well as a UWO MHSRB seed grant to AJM. Mass spectrometry work was supported by a research grant from the Keck Foundation, NIH grant R35 GM119536 and associated instrumentation supplement to JV. JRI and MDB were supported by an NSERC PGS-D and NSERC CGS-D respectively.

## Conflicts of Interest

The authors declare that there was no conflict of interest while conducting and reporting this research.

